# Spatial Proofreading Amplification of *in situ* Transcript and Protein Signals

**DOI:** 10.1101/2025.06.11.659176

**Authors:** Carsten H. Tischbirek, Katsuya L. Colón, Saori Lobbia, Christopher J. Cronin, Long Cai

**Author notes:** Corresponding author. **Materials & Correspondence** Material requests and correspondence should be addressed to Long Cai. These authors contributed equally (Carsten H. Tischbirek, Katsuya L. Colón).

## Abstract

Spatial transcriptomics experiments greatly benefit from brighter signals that improve detection efficiency and shorten imaging times. Here, we introduce a new enzyme-free signal amplification method inspired by the kinetic proofreading principle, in which short oligonucleotide probes are iteratively deposited at their target site and covalently photo-crosslinked to the cell, forming highly stable DNA assemblies. The method results in more than 500-fold signal amplification and supports multiplex readout capabilities.

## Main text

The kinetic proofreading principle, first introduced by Hopfield in 1974^1^ and Ninio in 1975^2^, describes how biological systems expend energy in sequential, irreversible checkpoints to lower the error rate of molecular recognition reactions, such as DNA replication or tRNA selection, beyond what equilibrium binding alone can achieve. We reasoned that this principle could be applied to *in situ* signal amplification applications, which are crucial for spatial omics methods to efficiently detect RNA transcripts and proteins. For example, current single molecule FISH (smFISH) methods^3–8^ can achieve 10,000-100,000 barcode multiplexing *in situ*, but suffer from low signal intensity levels and lack of robustness for fragmented RNAs. These methods would greatly benefit from brighter, higher quality signals with a low amplification error rate and low false discovery rate, which would also allow targeting of shorter RNAs and increased imaging speeds.

An ideal amplification method should satisfy three criteria. First, it should be highly efficient, i.e. every target detected by smFISH should yield an amplified signal that colocalizes with the smFISH signal, indicating that the underlying molecule is faithfully amplified. Second, amplification of real targets should be distinguishable from nonspecific amplification. Third, amplified signals should be compatible with sequential hybridization for multiplexing, with readily strippable and re-hybridizable signals that maintain a compact structure for the resolution of dense multiplexed signals. Existing amplification methods already excel at some of these criteria. Rolling-circle amplification (RCA)^9–12^, hybridization chain reaction (HCR)^13–15^, and branched-DNA (bDNA) approaches^16–18^, for example, show remarkable signal increases and versatility across different sample types. However, RCA’s enzymatically generated amplicons remain sparse, HCR produces non-compact DNA-structures and requires lengthy re-hybridization cycles, and the large DNA strands of bDNA can be limited by diffusion, significantly reducing their efficiency, and can non-specifically adhere to cells and tissue, generating off-target signals.

We designed SPARC (Spatial Proofreading Amplification with Recursive Crosslinking) to satisfy all these criteria. SPARC is based on the repeated, specific hybridization of an amplifier probe against its intended primary probe target, followed by an irreversible lock-in step that accumulates the amplifier at its correct target location as the amplification cycles are repeated, and a stable amplifier-seed is formed. This contrasts with off-target binding at incorrect locations, where amplifiers might bind nonspecifically, but are not accumulated over multiple rounds.

In detail, photoreactive benzophenone (BP)-tagged amplifiers^19–21^ (L1) first hybridize to a primary probe bound to cellular RNA. We next used UV-illumination to covalently anchor the hybridized amplifier to the cellular matrix (Figure 1Aa). A toehold strand then displaces the anchored amplifier from the primary probe (Figure 1Ab) to free up the initial amplifier binding site again (Figure 1Ac). Importantly, the displaced UV-crosslinked amplifier–displacer hybrid strand remains near its initial target yet cannot rebind to it (Figure 1Ad). Each cycle deposits a new amplifier at the same site, and multiple cycles assemble a stable amplifier seed structure that is localized on true targets (Figure 1Ae), whereas off target sites seldom accumulate crosslinked strands because every cycle starts over with the sequence-dependent hybridization step.

**Figure 1:**
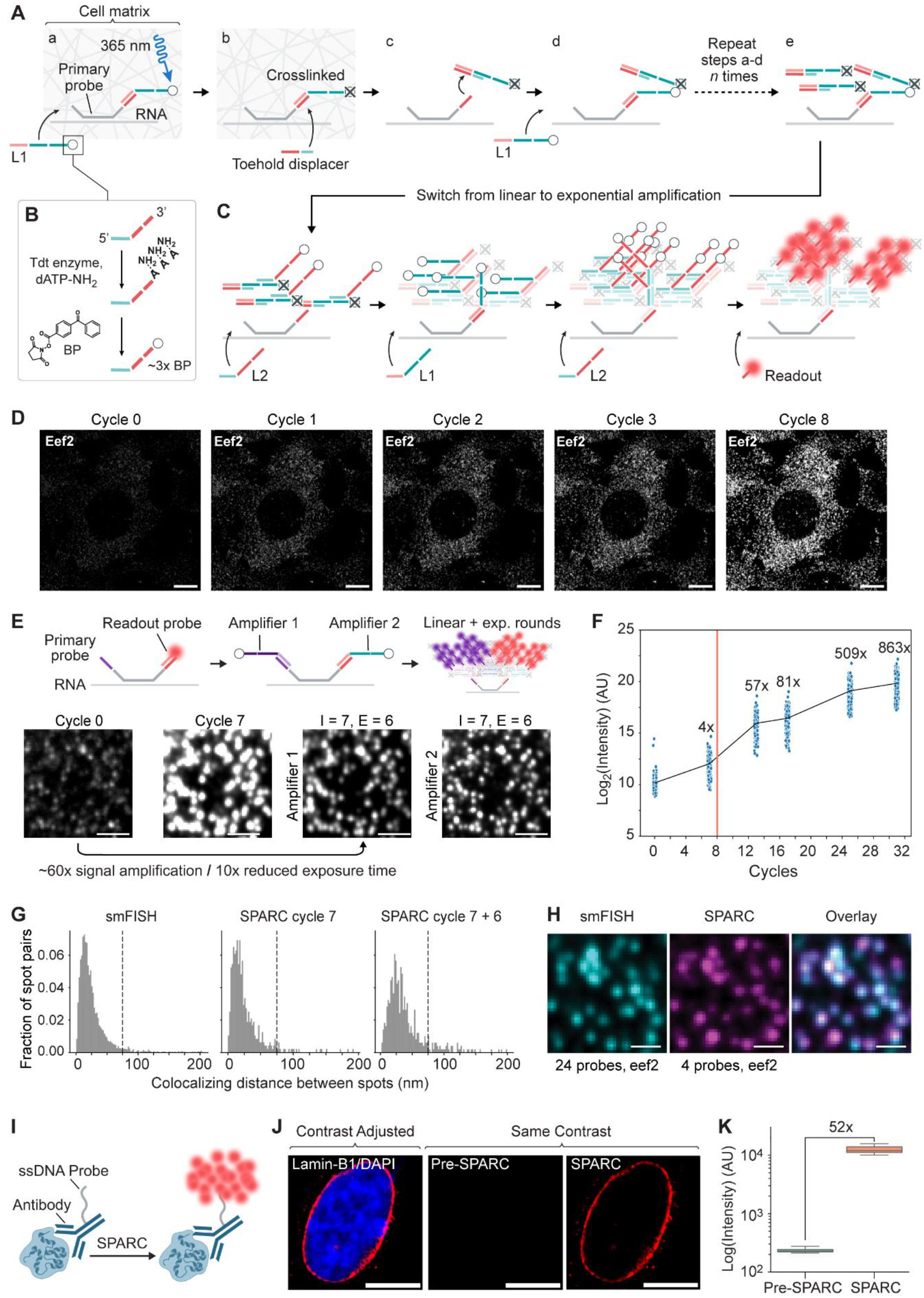
A) Schematic of SPARC amplification with (a) amplifier ‘L1’ hybridization and photocrosslinking, (b) displacement of photocrosslinked amplifiers, and (c) an unbound target probe for (d) subsequent amplification rounds to (e) crosslink multiple amplifier probes at the target site. B) Synthesis of SPARC amplifiers. C) Schematic of the exponential phase of SPARC amplification using an amplifier pair ‘L1’ and ‘L2’ and the addition of readout probes for imaging experiments. D) Representative images of linear signal amplification cycles as illustrated in panel A in NIH3T3 cells. Scale bar = 10 microns. E) Top: Schematic of the experiment to test combined linear and exponential amplification and colocalization of SPARC amplifier assemblies between two branches of the same primary probe. Bottom: Representative images of different amplification cycles. I = iteration, E = exponential, Scale bar = 2 microns. F) SPARC growth curve showing >500-fold signal amplification. Red line = start of exponential cycles. G) Representative distributions of the distance between two different SPARC generated branches on the same primary probe as illustrated in panel E, as well as smFISH (left panel). Dashed line = 75 nm. H) Representative images of NIH3T3 cells labeled with 4 primary probes for SPARC amplifiers and 24 probes for smFISH, and an overlay of both images illustrating their colocalization. Scale bar = 1 micron. I) Schematic of SPARC amplification of oligo-conjugated antibody signals. J) Left panel: Representative high contrast immunolabeling image of the nuclear lamina with Lamin-B1 antibody in HeLa cells recorded before SPARC amplification. Middle and right panels: The same cell recorded with short exposure times before and after SPARC amplification. Scale bar = 10 microns. K) Quantification of the signal increase in the representative experiment shown in Panel J. Horizontal line = median.

Photocrosslinkable amplifiers were prepared by terminal deoxynucleotidyl transferase extension with amine-modified dATP followed by reaction with benzophenone succinimidyl ester (BP), which attached about three BPs per strand (Figure 1B). To test the crosslinking efficiency, we designed BP-amplifiers with a USER cleavage site and a displacement toehold (Supplementary Figure 1A). After *in situ* USER digestion and harsh denaturing washes, almost all crosslinked strands remained, showing strong attachment of the amplifiers to the cell matrix. To test the efficiency of our displacer probe design with 10 nt toehold sequences, we applied displacers to non-crosslinked amplifiers and observed a clear reduction in signal intensity (Supplementary Figure 1B). Furthermore, we observed that displacer probes were required for amplifiers to bind the target probe in subsequent imaging rounds (Supplementary Figure 1B–1C).

Following linear amplification, additional assembly rounds on the iteratively formed seed can be performed without adding a displacer strand. Without a displacer strand, two secondary amplifier binding sites remain exposed, so that two amplifiers ‘L2’ can bind to the amplifier ‘L1’ and vice versa, allowing for exponential signal growth when switching between amplifiers ‘L1’ and ‘L2’ (Figure 1C) in subsequent rounds.

To test the SPARC concept, we imaged Eef2 RNA in NIH3T3 cells. Each cycle of SPARC amplification raised the signal intensity roughly linearly through multiple cycles (Figure 1D). The signal remained colocalized with smFISH spots during the amplification rounds (Figure 1E, ‘Cycle 0’). Six additional exponential rounds increased the signal by about 60-fold total while each amplified spot stayed diffraction limited and still colocalized with the smFISH signal. Additionally, two branches that grew simultaneously from one primary probe colocalized with each other, which indicated uniform growth across branches (Figure 1E, right panels). This was consistent with a simulation showing a reduced overall variance in exponential amplifier assembly across both branches when a linearly amplified seed was used (Supplementary Figure 2). Extended scaffold growth pushed the total signal gain beyond 500-fold (Figure 1F), although it is noteworthy that later rounds yielded less signal increase per amplification round compared to a theoretical exponential growth curve, likely due to amplifier depletion, dye self-quenching, or saturated crosslink sites at very high amplification factors.

Localizing the centroid of the SPARC dots showed that two separate SPARC branches, amplified independently from the same RNA with 7 linear and 6 exponential cycles and around 60-fold signal increase, remained within 50 nm of each other, which was similar to smFISH (Figure 1G, Supplementary figure 3). Importantly, with these amplification parameters, four primary probes targeting Eef2 RNA yielded amplified spots that colocalized well with 24 smFISH primary probes (∼70% colocalization, Figure 1H). We were therefore able to efficiently detect a low number of primary probe targets, which allows imaging of shorter RNA targets and lowers experimental costs and complexity.

To test the applicability of SPARC beyond the detection of RNA targets, we also amplified antibody signals (Figure 1I). In HeLa cells, the signal of a Lamin B1 antibody conjugated to an 18 nt DNA oligonucleotide was amplified over 50-fold while retaining the nuclear lamina pattern that matched unamplified controls. Here, SPARC amplification was able to achieve a dense and continuous immunolabeling pattern that matched a non-amplified antibody staining (Figure 1J-K), which can be challenging for sparse amplification methods relying on enzymatically generated DNA amplicons. The method therefore extends to low abundance proteins and hard to detect antibody signals, with the advantages of lowering the required antibody concentration for successful stainings and that the resulting DNA structures can withstand repeated denaturing washes, which allows multiplex fluorescent signal readout.

To examine multiplex capacity with multiple amplifiers simultaneously, we applied SPARC to 16 RNA species in NIH3T3 cells (Figure 2A). Individual amplifiers can vary in amplification factors, spanning 30 to 70-folds (Figure 2B). Amplifiers are colocalized for all 16 genes that were imaged (Figure 2C), but had varying background nonspecific-binding levels (Supplementary Figure 4). In multiplex experiments, we observed that high total amplifier concentrations beyond 1 µM can lead to amplifier aggregate formation in samples, which was not observed with BP-free amine control oligos (Supplementary Figure 5). Optimizing hybridization buffers for high amplifier concentrations, we observed that neutral crowding agents such as polyethylene glycol induced liquid-like condensates at high concentrations, whereas anionic dextran sulfate promoted robust binding without condensation (Supplementary Figure 6).

**Figure 2:**
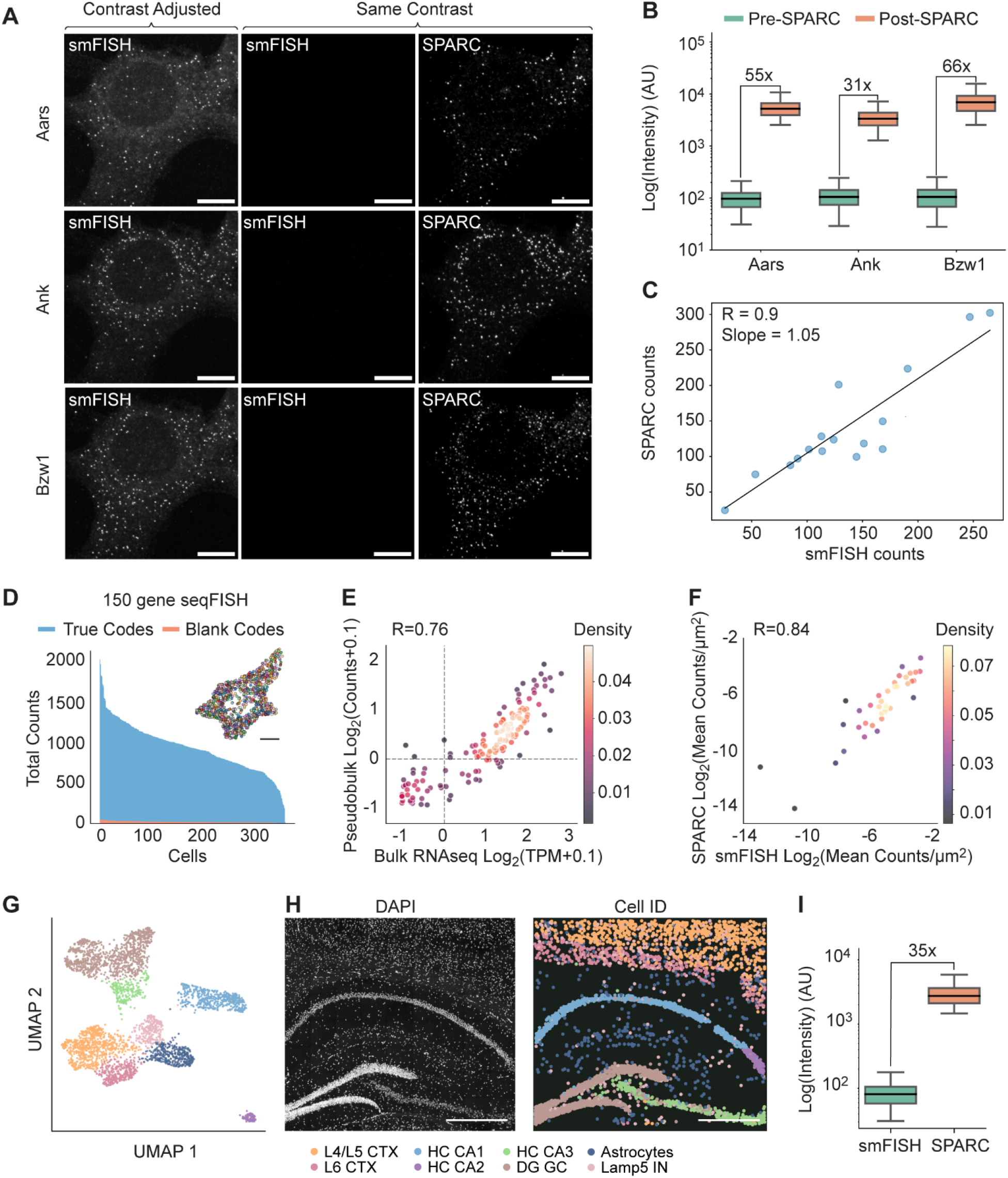
A) Representative images of different mRNAs in NIH3T3 cells imaged with smFISH and SPARC. Scale bar = 10 microns. B) Quantification of smFISH and SPARC images showing amplification factors for three example mRNAs shown in A. Horizontal line = median. C) Regression between SPARC generated structures and smFISH of the same RNA targets from the same cell shows high correlation. D) Counts distribution of a 150-plex SPARC in NIH3T3 cells experiments. Insert above distribution: Example 3T3 cell with detected RNAs color coded for different RNA species. Scale bar = 10 microns. E) Correlation of SPARC-amplified seqFISH RNA counts with bulk RNA-seq counts. F) Correlation of SPARC-amplified seqFISH RNA counts with smFISH counts. G) UMAP of SPARC-amplified 161-gene experiment in a mouse coronal brain slice. Cell types color-coded as listed in panel H. H) Representative image showing a DAPI-stained mouse coronal brain slice (left) and cell-type map (right). Cell type annotation shown below. Scale bar = 0.5 mm. I) Quantification showing 35-fold signal amplification obtained from colocalizing smFISH and SPARC-amplified spots in the representative dataset shown in G-H before and after amplification. Horizontal line = median.

The high stability of SPARC scaffolds under denaturing washes enabled a 150-plex seqFISH SPARC experiment in NIH3T3 cells using six pseudocolors, three barcode rounds, and one parity round. We decoded on average 913 ± 287 (s.d.) RNA counts per cell (n = 370 cells, see also Supplementary Figure 7) and observed a strong agreement between smFISH and RNA sequencing (Figure 2D-F) with a 5.0% FDR. Furthermore, to test the concept of SPARC in tissue slices, we used the same encoding scheme as in the cell culture experiments to profile 161 genes in coronal mouse brain slices. We recovered known cell classes in the hippocampal area including CA1, CA2, CA3, dentate gyrus, and cortical layers five and six (Figure 2H), and observed more than 30-fold signal amplification with SPARC in intact mouse brain slices (Figure 2I), markedly reducing the imaging time required for the sample compared to non-amplified samples.

In summary, we show that the concept of spatial proofreading used for SPARC amplification allows for the generation of high efficiency, low background, stable and compact DNA-assemblies that boost signal up to 500-fold while maintaining high spatial accuracy. Future work will involve screening additional amplifiers to further improve detection efficiency and specificity, as well as extending the method to more complex biological specimens including multi-omics imaging experiments.

## Acknowledgements

We thank Inna-Marie Strazhnik for help with figures, Jina Yun for excellent technical support, and the Caltech Proteome Exploration Laboratory for access to laboratory equipment. This project was supported by NIH DP1 NS131408.

## Author Contributions

C.H.T., K.L.C. and L.C. conceived the idea and designed experiments. C.H.T., K.L.C., S.L. and C.J.C. prepared and validated experimental materials. C.H.T., K.L.C. and S.L. performed experiments. C.H.T., K.L.C. and L.C. performed data analysis, made the figures and wrote the manuscript with input from all authors. L.C. supervised the project.

## Competing interests

C.H.T., K.L.C. and L.C. filed a patent application on the SPARC amplification. L.C. is co-founder of Spatial Genomics Inc.

